# Functional Brain Connectivity and Musical Training: Insights from EEG Phase Synchronization During Music Perception

**DOI:** 10.1101/2024.12.11.627938

**Authors:** Anna A. Provorova, Sofya P. Kulikova

**Author notes:** corresponding author: Sofya Kulikova, HSE University, blvd. Gagarina, 37a-218, Perm, Russia.

## Abstract

Musical training profoundly impacts brain organization, fostering functional and structural adaptations that enhance auditory, motor, and cognitive processes. This study investigates functional connectivity differences between professional musicians and non-musicians during tasks involving music perception and different graphic notations. Using EEG phase-locking value (PLV) analysis, we examined synchronization across alpha, beta, and gamma frequency bands. Results revealed that musicians exhibited enhanced functional connectivity between brain regions, reflecting their ability to integrate auditory, visual, and cognitive information efficiently. In the alpha band, strong connectivity between the occipital and temporal regions was observed during all tasks, with additional synchronization between the parietal and temporal regions in tasks lacking pitch information. In the beta band, significant frontal-temporal connectivity was detected, indicating cognitive processing, while temporal-occipital synchronization emerged in specific tasks. In the gamma band, musicians demonstrated robust frontal-temporal and frontal-frontal synchronization, reflecting emotional and structural processing of music. Non-musicians showed reduced task-specific connectivity, relying on generalized strategies. These findings underscore the role of musical training in enhancing multimodal brain integration and cognitive flexibility, contributing to advanced music perception and processing.

## Introduction

Musical training has a profound effect on brain organization and functioning. Conducting post-mortem studies more than one hundred years ago Auerbach (1906) had noticed that brain temporal lobes of famous musicians were more developed than in other people, who did not practice music. With the invention of non-invasive neuroimaging techniques, like MRI (Magnetic Resonance Imaging), it became obvious that the brains of musicians have important structural differences compared to non-musicians. Such differences have been reported for the auditory (Schneider et al., 2002), somatosensory (Elbert et al., 1995) and motor (Schlaug, 2001) cortices, as well as for the corpus callosum (Lee et al., 2003) and the cerebellum (Hutchinson et al., 2003). These areas are directly involved in auditory processing, motor control, and cognitive functions and the increased cortical thickness there likely reflects the effects of neuroplasticity induced by extensive musical training. Indeed, it was shown that musicians have greater cortical thickness in the left superior frontal gyrus and deeper sulci in the auditory-related areas (Rus-Oswald et al., 2022; Liu et al., 2024), which may stem from an intensive activity of the neuronal network involved in mnemonic retention, monitoring, and retrieval (Bermudez et al., 2009) during musical training. At the same time, the increased cortical volume observed in motor and visual areas reflects the enhanced spatial-temporal attention and sensorimotor transformation (Alghamdi, 2023). Furthermore, structural covariation studies demonstrate that early trained musicians have larger volumes in brain areas that promote coordinated developmental plasticity between the cerebellum and cortex, which is required for successful integration of spatiotemporal and motor information (Shenker et al., 2022). However, morphological findings can only indicate possible effects that musical training exerts on brain functioning and they themselves say little about the processes that underlie music perception.

Functional and behavioural studies highlight the tremendous effect that musical training has on brain cognitive development. The neural mechanisms that differentiate musicians from non-musicians are likely to reflect functional adaptations, induced by musical training and experience. It has been demonstrated that musicians exhibit enhanced auditory processing capabilities, greater integration of acoustic features, and specialized neural networks that facilitate music perception and performance. In particular, by investigating the properties of the mismatch negativity response (MMN) using magnetoencephalography (MEG), Hansen et al. (2022) have shown that musicians have a more integrated neural processing of complex acoustic features, specifically for frequency-related deviants in complex, melody-like stimuli. Furthermore, the authors suggested that this enhanced ability to process intricate musical elements arises from the recruiting of the overlapping neural resources for contextually relevant stimuli. The results of a recent fMRI study by Jiang et al. (2022) suggest that musicians also show a superior processing of musical syntax thanks to a greater involvement of the dorsal stream, including the inferior frontal and superior temporal gyri, which is crucial for tonality perception. Thus, musical training enhances sound sensitivity, verbal abilities, and reasoning skills, contributing to the overall cognitive development (Miendlarzewska & Trost, 2014).

The contribution of musical experience to brain development is not limited to the promotion of auditory-related skills. Musical experience also aids in emotional regulation and creativity, and is essential for holistic brain development (Yazar, 2024). Numerous studies indicate that music training has a beneficial effect on both overall academic performance and the development of individual cognitive skills, such as speech comprehension and production (Patel, 2011), spatial-temporal problem solving (Rauscher et al., 1997), fractional arithmetic (Azaryahu et al., 2020), etc. Such a positive effect is likely based on universal learning mechanisms activated in common and/or adjacent areas of the brain (Patel, 2011). However, music learning, and in particular, mastering musical notation, is not simpler than learning a language or mathematics, as it requires effective multimodal integration of auditory, visual, and motor processes. Disruption of such integrative processes can cause difficulties both in music learning and in general education.

At the neuronal level, the effectiveness of the integration processes may be evaluated using such techniques as EEG (electroencephalography), fMRI or MEG. For example, the degree of synchronization of the brain electrical oscillations, recorded using EEG from different brain areas, can serve as a measure of the effectiveness of such integrative processes (Picazio et al., 2024; Jerde et al., 2011). Indeed, in the work of Picazio et al. (2024) it has been indicated that the simultaneous perception of the pitch and rhythm of a melody requires the coordinated work of the frontal-cerebellar network, and interference with its work leads to a disruption of music perception. The study of Jerde et al. (2011) also suggests that the melody and its rhythm are analyzed by different areas of the brain, which must be synchronized with each other. At the same time, the effectiveness of synchronization depends on musical experience and differs between professional and amateur musicians (Papadaki et al., 2023; Pallesen et al., 2015). Thus, measuring synchronization may present a potential tool for assessing and monitoring the acquisition of music perception skills.

Indeed several studies suggest that musical training leads to enhanced intra- and inter-network functional connectivity, which is associated with the improved efficiency and automaticity in brain functions (Hou et al., 2024). Experienced musicians demonstrate stronger connections and higher efficiency in the brain networks, linked to auditory tasks such as interval recognition (Papadaki et al., 2023), and have stronger integration between sensory, motor, and cognitive networks. In contrast, non-musicians rely more on default-mode networks during music processing (Alluri et al., 2017) and in particular, on visual cues during audiovisual integration (Paraskevopoulos et al., 2015), indicating a more generalized approach to music perception. This highlights the profound impact of musical training on brain functioning, suggesting that the differences in neural mechanisms are not merely quantitative but qualitative in nature (Niranjan et al., 2019).

The connectivity patterns in musicians may vary significantly based on their level of expertise, revealing distinct neural adaptations associated with musical training. For example, musicians with improvisational training exhibit different inter-network connectivity compared to classically trained musicians, indicating a more dynamic neural response to musical stimuli, which was revealed in alpha (8-12 Hz) and beta (15-30 Hz) ranges of the EEG oscillations (Faller et al., 2021). All together, these findings suggest that musical training enhances specific connectivity patterns, yet it is important to keep in mind that not all musicians may equally express such connectivity patterns, suggesting variability based on individual training and experience levels (Leipold et al., 2021). Taking into account the important role of the brain synchronization in successful acquisition of the musical skills, the aim of this work was to deepen the understanding of its contribution to the process of musical notation perception, and in particular, to determine those functional connections that are responsible for the integration of visual, frequency (pitch) and rhythmic (beat) information. To achieve this, we have conducted an experimental study using a specially developed paradigm that involves investigating functional connectivity based on EEG signals analysis during the process of simultaneous listening to musical melodies and reading their musical notation. The experimental design and the results of the study are described in the following sections.

## Methods

### Participants

The experiment involved 47 healthy participants, including 29 women and 15 men, with a mean age of 30 years (SD = 11.3). Among these participants, 34 were assigned to the musicians’ group, while the remaining individuals constituted the control group. All members of the musicians’ group had classical musical education for at least 8 years at a music school, and they continue to play musical instruments. The average experience of musicians in this group was 21 years (SD = 12.9). Participants in the musicians’ group had received training on a variety of musical instruments, including piano, guitar, violin, flute, accordion, bayan, and balalaika. In contrast, individuals in the control group had no prior training in playing musical instruments or in music theory. The selection criteria ensured that all musicians were actively engaged in music practice, while the control group participants had no previous exposure to musical training. The demographic characteristics of the participants were balanced to minimize confounding variables, thereby enhancing the validity of the findings. All participants have signed the informed consent and the study was approved by a local ethics committee.

### Experimental Tasks

During the experimental session, all respondents completed a specific set of tasks designed to evaluate their skills in music notation perception. Participants were asked to match a short musical excerpt they listened to with its symbolic representation, reflecting various aspects of musical notation, and to identify the last note played. The musical stimuli consisted of monophonic exercises of varying complexity drawn from classical solfeggio textbooks. The graphical representation of the melodies could take one of the following forms (see Figure 1):

- Type 1: A “classical” notation;
- Type 2: A representation containing only the rhythmic pattern without pitch information;
- Type 3: A representation containing information solely about pitch;
- Type 4: A representation indicating only the number of notes in the melody;
- Type 5: A representation containing verbal description of the melody. The notes are expressed in words, while the rhythmic information is absent.

**Figure 1.**
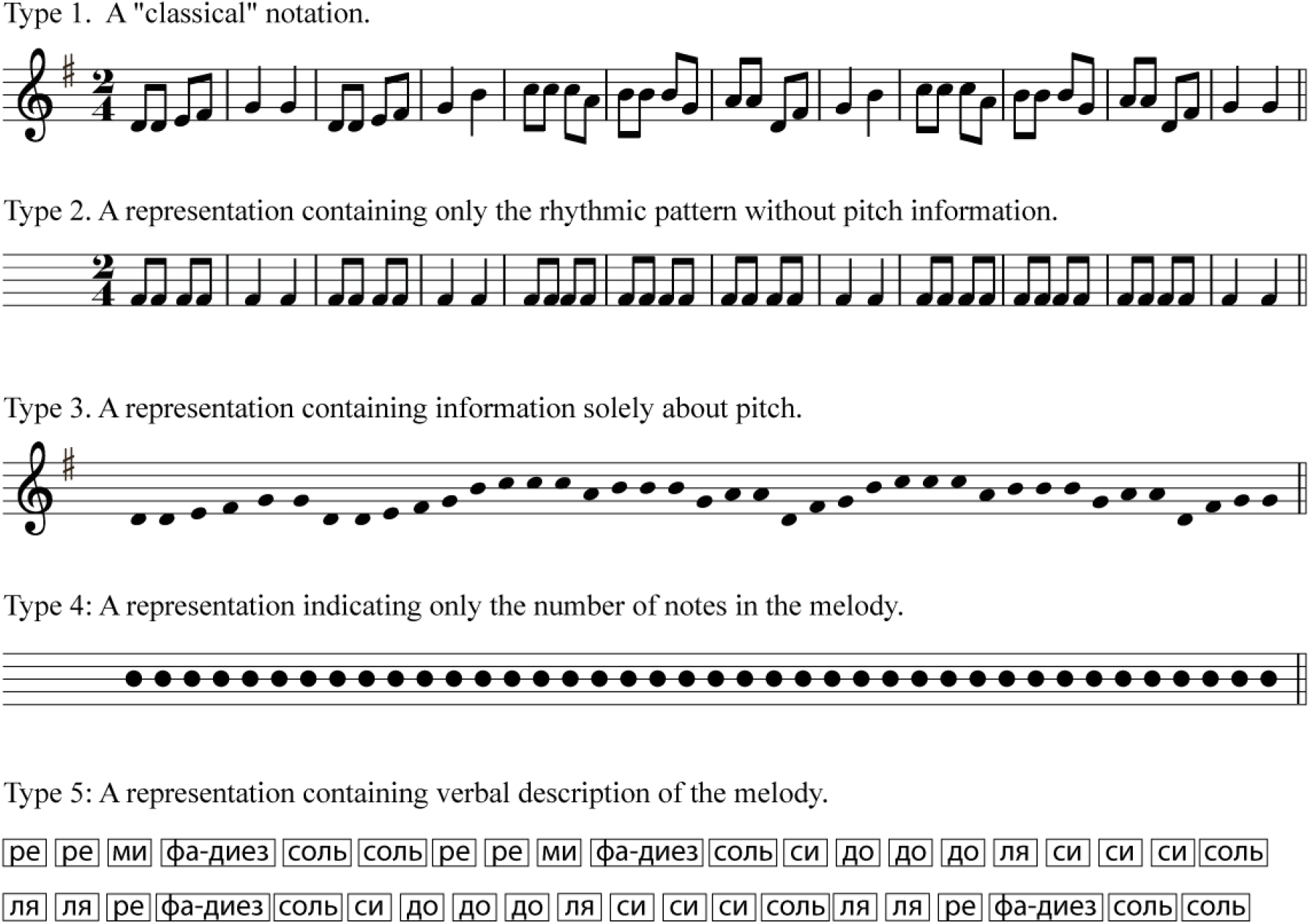
Examples of graphical representations of melodies in the tasks.

Each participant was presented with audio stimuli in a random order, and the corresponding graphical representation was also randomly selected from the five aforementioned variants. The stimuli were delivered using the in-house software “NoteIT” (available on request), developed by authors in HTML, CSS, JavaScript utilizing the jsPsych library (Leeuw et al., 2023). This software recorded timestamps and the selected note for each trial allowing to estimate the reaction time and the accuracy of the answer. The demonstration of the tasks to the participants was conducted using a web browser (Chrome 130.0.6723.117). There was no time limit for responses in each task; however, the reaction time was also recorded. Overall, the study included a total of 30 different melodies with varying durations (ranging from 6-8 notes to 40 or more), played in both fast and slow tempos.

### EEG acquisition

During the experimental session, participants were sitting in comfortable chairs in front of a 22-inch computer screen, which was positioned approximately 80 cm from their eyes. The sound stimuli (monophonic) were presented through two speakers positioned 80 cm in front of the participants. The recording took place in a quiet room, and prior to the recording, participants were instructed that any extraneous movements or sounds could adversely affect the quality of the EEG recording. Therefore, they were asked to minimize their movements during the listening tasks and to refrain from singing the melodies aloud.

EEG data from participants in the experiment were collected using a wireless electroencephalograph “NeuroPolygraph” (Neurotech, Russia). This electroencephalograph features 21 channels arranged according to the 10-20 system at FP1, FPZ, FP2, F3, FZ, F4, F7, F8, C3, CZ, C4, P3, PZ, P4, T3, T4, T5, T6, O1, OZ, O2, using carefully positioned nylon caps, with reference electrodes placed on the mastoid processes behind the ears. Impedance was kept below 20 kΩ. The sampling frequency was 500 Hz. Data was transmitted via Bluetooth to a laptop, allowing the experiment organizer to monitor the progress of the experiment in real-time and to save the data using software provided by Neurotech.

EEG recordings were obtained during the completion of the tasks described in the previous section. In addition, data on blink frequency were collected using an electrode attached to the left eyelid, which facilitated the subsequent removal of blink artifacts from the data. Prior to the commencement of the tasks, a baseline period of 20 seconds of silence was recorded, after which the participants began the testing phase. Participants had the opportunity to rest after each exercise, as intermission screens without any information were presented between tasks. A “Next” button was available for participants to click when they were ready to proceed with the test.

### EEG pre-processing

The processing of EEG data was conducted offline using Python 3.10, specifically utilizing the MNE library (Gramfort et al., 2013). This library provides a comprehensive suite of tools for the analysis of electrophysiological data, enabling the efficient handling and interpretation of the recorded EEG signals. The offline processing included steps such as filtering, segmentation and artifact rejection of the data to prepare it for further analysis.

The EEG data for each participant were filtered with a lower cutoff frequency of 0.5 Hz (−6 dB cutoff 0.25 Hz) and an upper cutoff frequency of 50 Hz (−6 dB cutoff: 56.25 Hz). This filtering range was chosen to eliminate low-frequency noise, such as drift and movement artifacts, while preserving the relevant brain activity signals typically found within this frequency band.

Subsequently, the data were first splitted into epochs and only then subjected to artifact cleaning, because this improves the quality of the data cleaning process by allowing for a more focused analysis of the segments of interest. Timestamps of events marking the beginning of melody listening were recorded using a specialized software that were used as the beginning of epoch, and the durations of all melodies were known. Consequently, each epoch consisted of the listening process of a melody until its end, with an additional 2 seconds added to both ends to mitigate edge effects that may arise during analysis.

Then the data were cleaned of artifacts using Independent Component Analysis (ICA) (Hyvärinen & Oja, 1997), specifically employing the infomax algorithm. Since the electrooculogram (EOG) signal was also recorded during the EEG data collection, components that were most correlated with EOG were selected and removed to address blink artifacts.

Next, to eliminate the possibility of the volume conduction phenomenon, a transition to a Laplacian reference was performed. This approach is implemented to reduce the effects of distant activity on the recorded signals, thereby enhancing the spatial resolution of the EEG data and allowing for a more accurate localization of the neural sources underlying the observed potentials.

Finally, before estimating the synchronization between EEG channels, the selected segments from each EEG data were processed with a zero phase distortion finite impulse response (FIR) filter (filter order: 256) across three frequency bands (FBs): alpha-α (8–13 Hz), beta-β (13–30 Hz), and gamma-γ (30–50 Hz).

### Data analysis

To evaluate the functional connectivity (FC) between pairs of EEG channels, we employed the phase-locking value (PLV) (Bruña et al., 2018), a phase-synchronization (PS) index, which is a measure of the consistency of phase differences between two signals over time. It quantifies the degree to which the phases of two signals are aligned, providing an insight into the synchrony of neural oscillations. To compute the PLV between two noisy real-valued signals *x*_*k*_(*t*) and *x*_*l*_ (*t*), the following steps are undertaken. First, the phases of each signal are estimated by constructing analytical signals of two narrowband signals using the Morlet wavelet transform. Once the signals are transformed, the instantaneous phases ϕ_*k*_(*t*) and ϕ_*l*_(*t*) for each time point t are obtained. Then the relative phase difference is calculated. The next step involves constructing a complex exponential from the relative phase as 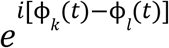 where *i* is the imaginary unit. The PLV is calculated by averaging the modulus of these complex exponentials over the entire time series:

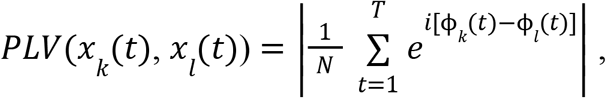

where *N* is the total number of timepoints. The resulting value is often normalized to provide a measure between 0 and 1, where 0 indicates no PS and 1 indicates perfect PS. This calculation was implemented using the spectral_connectivity_time function from the MNE software package, which facilitates the analysis of functional connectivity in neurophysiological data.

First, for each participant, the authors calculated the phase-locking value (PLV) matrices during a 20-second period of silence recorded before the onset of the experiment. This was done to examine the functional connections observed in the participants’ brains during inactivity. Subsequently, PLV matrices were computed separately for five different types of tasks, with data within each type obtained by averaging the results from six tasks of the same type.

From the resulting matrices for each task type, the data regarding connectivity during inactivity were subtracted. In cases where a cell produced a value less than zero, it was replaced with zero. Then the data on phase synchronization between all channels (a total of 210 pairs from 21 electrodes) for both the musicians’ groups and the control group were pairwise tested for statistical differences using the Mann-Whitney U test. Only those connections that exhibited statistically significant differences at the 5% level were selected. After processing the data in this manner for each respondent, the resulting matrices for musicians and the control group were averaged, in order to obtain overall assessments of phase synchronization across the different task types and only statistically significant results are shown.

The data about the accuracy of responses and reaction times among musicians and the control group were also analyzed. To compare the samples, the Mann-Whitney U test was employed. To compare the results by type for the control group, the Wilcoxon test was employed, and the analysis was conducted using Python 3.10.

## Results

The accuracy of responses across almost all types of tasks among musicians significantly differed from that of the control group, favoring more precise answers (see Figure 2). While participants in the control group had approximately the same number of wrong notes across various tasks and the difference by types is not statistically significant (Type 1: mean = 3.14 notes, Type 2: mean = 3.42 notes, Type 3: mean = 4.67 notes, Type 4: mean = 3.6 notes, Type 5: mean = 3.78 notes), musicians placed considerable importance on the form of musical notation. In tasks of Type 1 (classical notation) and Type 3 (only pitch information), musicians exhibited nearly no errors in their responses (Type 1 mean = 0.9 notes, Type 3 mean = 0.8 notes). However, in tasks lacking pitch information, the number of errors increased. In task Type 4 (only indicated the number of notes), the results of musicians and the control group were statistically indistinguishable (p-value = 0.14).

**Figure 2.**
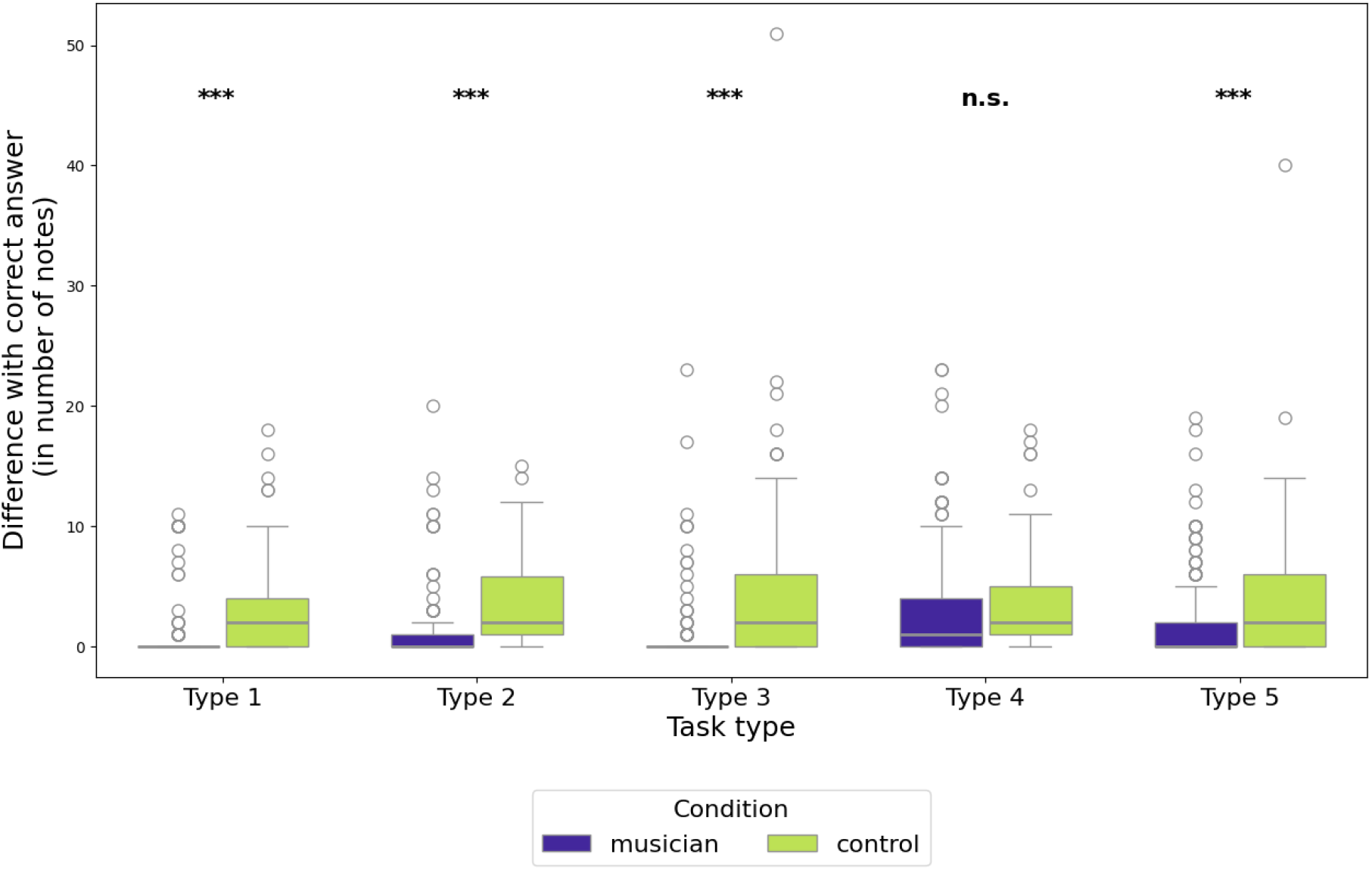
Type-by-type comparison of difference with the correct answer between musicians and control group, *** - p-value < 0.01, ** - p-value < 0.05, * - p-value < 0.1, n.s. - not significant

The reaction time before responding in the control group was significantly shorter in some types of tasks compared to musicians (see Figure 3). In tasks without pitch information (Types 2 and 4), musicians required significantly more time to contemplate their responses than the control group. Additionally, in task Type 5, musicians also needed significantly more time for reflection. Conversely, participants in the control group spent approximately the same amount of time considering their responses, regardless of the type of task and no statistically significant differences were found between their results by types (Type 1: mean = 1.99 sec, Type 2: mean = 2.32 sec, Type 3: mean = 2.36 sec, Type 4: mean = 2.53 sec, Type 5: mean = 3.11 sec).

**Figure 3.**
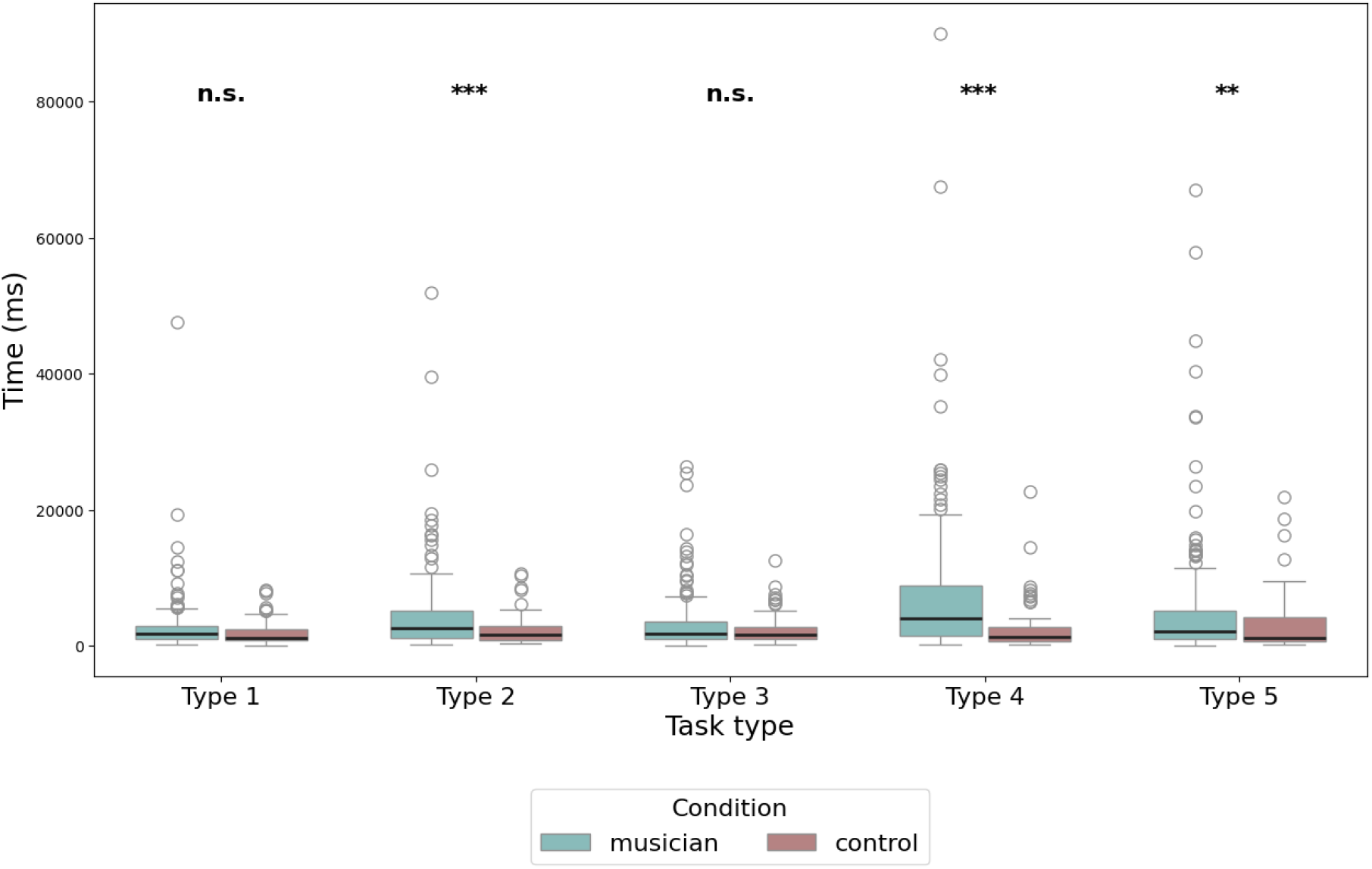
Type-by-type comparison of reaction time between musicians and control group, *** - p-value < 0.01, ** - p-value < 0.05, * - p-value < 0.1, n.s. - not significant

Furthermore, no statistically significant differences in accuracy and response speed between male and female participants within the musicians’ group were found.

Figure 4 shows the top 10 statistically significant functional (phase) connections between various regions in the alpha, beta, and gamma bands. Values above 0 indicate connections that are clearly expressed in musicians, while values below 0 represent connections that are more expressed in the control group. There were no statistically significant functional connections characteristic of the control group that were absent in musicians at a phase locking value (PLV) level above 0.05.

**Figure 4.**
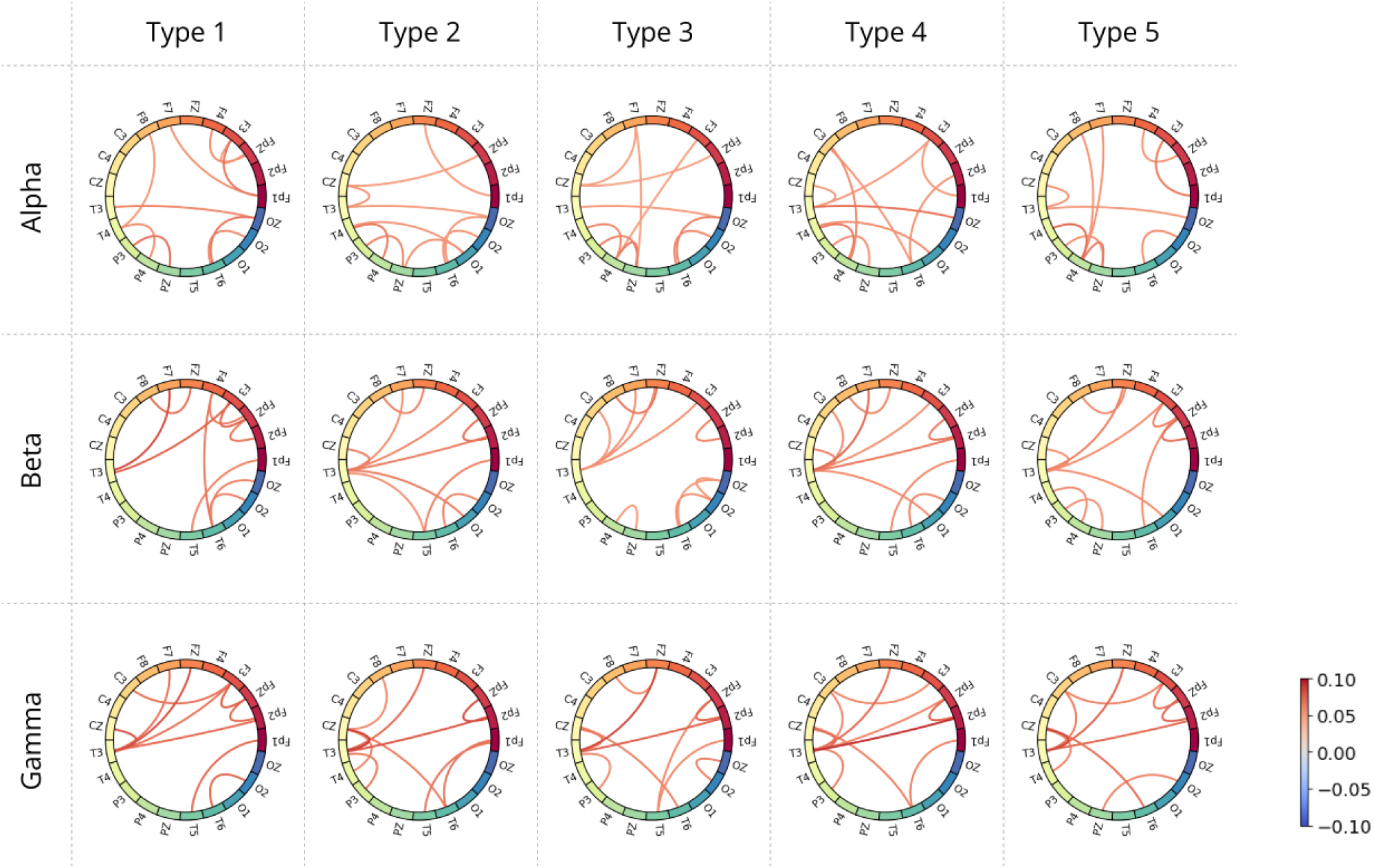
Functional (phase) connectivity between pairs of EEG leads in alpha, beta and gamma bands.

## Discussion

The comparison between musicians and non-musicians reveals some intriguing differences in brain activity during music listening. Research indicates that musicians exhibit increased phase synchrony in alpha, beta and gamma bands, suggesting they have stronger functional connectivity in these frequency ranges. This enhanced synchrony may be linked to their extensive training and experience in music, allowing them to process musical elements more efficiently. This finding aligns with Eierud and colleagues’ (2023) demonstration that lifelong musicianship may contribute to enhanced brain and cognitive reserve.

Furthermore, there are notable hemispheric differences in brain activity: musicians tend to exhibit stronger synchrony in the left hemisphere during music perception. These finding may indicate that musicians utilize more language-related areas of the brain when processing music, as the left hemisphere is traditionally associated with language and analytical tasks. This could imply that musicians are not only hearing music but also interpreting it in a more structured and nuanced way, akin to how they would process language. Another explanation for the asymmetry in brain hemisphere activity during task performance may be related to the idea proposed by Rosenthal (2016) that the lateralization of expectations can vary based on the type of expectation generated by the context. Specifically, higher-level processing is predominantly managed by the left hemisphere for sequentially ordered expectations, while the right hemisphere is more active for non-sequential expectations. The tasks presented in this study can indeed be classified as involving sequentially presented expectations, as musicians frequently noted that while reading the graphical representation of a melody, they would glance ahead to the note currently being played in order to “anticipate” the music they were hearing.

Next, we will analyze the results separately across different frequency bands. In the alpha band, a strong connection was identified between the occipital lobe (Oz) and the temporal lobe (T3), regardless of the type of task performed. However, in tasks where the representation of notes lacks information about pitch, a statistically significant enhancement of the functional connection between the parietal lobe (Cz) and the temporal lobe (T3) was observed, which may indicate the integration of auditory information into the cognitive process.

In the beta band, a strong connection between the frontal regions of the brain (F7, Fz, F3) and the left temporal lobe (T3) was also noted, irrespective of the type of task performed. This may suggest significant cognitive processing occurring during these tasks. The strong connectivity in the beta range implies that there is an integrative network at play, where cognitive control mechanisms in the frontal lobe closely interact with auditory processing in the temporal lobe. Furthermore, in tasks that lack pitch information (types 2, 4, and 5), a statistically significant synchronization between the temporal cortex (T3) and the occipital cortex (O1) emerges, in contrast to the non-musician group.

In the gamma band, synchronization in the frontal region between FpZ and Fp2 was observed across all tasks, indicating that musicians actively process complex musical structures. This is a clear distinction from the non-musician group, which demonstrates statistically indistinguishable results regardless of the type of task, suggesting that they process tasks uniformly, irrespective of the representation of the musical melody. Indeed, participants in the control group are entirely unfamiliar with the graphical representation of music and cannot read it; thus, their task performance, regardless of type, was reduced to counting notes. Additionally, it is noteworthy that in the gamma band, musicians exhibited strong synchronization between the frontal lobe (Fp2, Fz) and the temporal region (T3) across all task types, further emphasizing the significant involvement of auditory information in task resolution. The level of synchronization was highest in the gamma frequency range compared to other ranges, aligning with the findings of Bhattacharya and Petsche (2000). Their research demonstrated that musicians show a significantly greater degree of phase synchrony in the gamma range while listening to music, in contrast to non-musicians. This indicates that musical training may improve the brain’s capacity to synchronize activity in this frequency range during the processing of music.

The results regarding the reaction time to tasks by musicians and non-musicians doesn’t align with previously published findings by Medina & Barraza (2019), which indicate that the reaction time of individuals engaged in music is statistically significantly shorter than that of non-musicians. For tasks that are of a specific nature, involving reading specific musical notation, listening to music, and comparing the two, we can state that musicians respond more slowly than non-musicians. This is likely related to the fact that musicians experience a greater cognitive load; while listening to music and extracting information from the specific graphical representation of notes, they are more inclined to strive for an accurate response. Conversely, non-musicians exhibit considerably less motivation to respond correctly, as their expectations of themselves are inherently lower, and according to Rosenthal (2016), expectations indeed play a crucial role in how listeners perceive music.

## Conclusion

This study highlights the profound impact of musical training on functional brain connectivity, particularly in the context of music perception and the integration of multimodal information. By analyzing EEG phase synchronization across different frequency bands during tasks involving musical notation perception, we demonstrated significant differences between musicians and non-musicians.

Musicians exhibited enhanced functional connectivity across alpha, beta, and gamma frequency bands, with notable synchronizations between frontal, temporal, parietal, and occipital brain regions. These findings suggest that musicians possess a more efficient and specialized neural network for processing complex auditory and visual information. In contrast, non-musicians lacked task-specific synchronization patterns and processed musical stimuli in a more uniform and generalized manner, relying primarily on visual cues and simpler cognitive strategies. This difference likely reflects their unfamiliarity with musical notation and the absence of specialized neural adaptations associated with musical training.

These results provide new insights into the neural mechanisms underlying music perception and the role of musical training in shaping functional brain connectivity. The data emphasize the importance of multimodal integration and hemispheric specialization in musicians, reflecting their ability to process complex musical stimuli efficiently. Furthermore, this study highlights the potential of EEG-based connectivity analysis as a tool for evaluating and monitoring the development of musical skills and auditory-cognitive integration. Future research could explore how different levels and types of musical expertise influence connectivity patterns and whether similar enhancements can be observed in other domains of learning that require multimodal integration.

## Acknowledgment

This article is an output of a research project implemented as part of the Basic Research Program at the National Research University Higher School of Economics (HSE University).

